# High proportions of single-nucleotide variations associated with multidrug resistance in swine gut microbial populations

**DOI:** 10.1101/2022.12.03.518979

**Authors:** Brandi Feehan, Qinghong Ran, Kourtney Monk, T. G. Nagaraja, M. D. Tokach, Raghavendra G. Amachawadi, Sonny T M Lee

## Abstract

**Background:** Antimicrobial resistance (AMR) is a significant global public health concern associated with millions of deaths annually. Agriculture has been attributed as a leading factor in AMR and multidrug resistance (MDR) associated with swine production estimated as one of the largest agricultural consumers of antibiotics. Therefore, studying and understanding AMR in swine has global relevance. AMR research has received increased attention in recent years. However, we are still building our understanding of genetic variation within a complex gut microbiome system that impacts AMR and MDR. In order to evaluate the gut resistome, we evaluated genetic variation before, during, and after antibiotic treatments. We studied three treatment groups: non-antibiotic controls (C), chlortetracycline (CTC) treated, and tiamulin (TMU) treated. We collected fecal samples from each group and performed metagenomic sequencing for a longitudinal analysis of genetic variation and functions.

**Results:** We generated 772,688,506 reads and 81 metagenome assembled genomes (MAGs). Interestingly, we identified a subset of 11 MAGs with sustained detection and high sustained entropy (SDHSE). Entropy described genetic variation throughout the MAG. Our SDHSE MAGs were considered MDR as they were identified prior to, throughout, and after CTC and TMU treatments as well as in the C piglets. SDHSE MAGs were especially concerning as they harbored relatively high variation. Consistently high variation indicated that these microbial populations may contain hypermutable elements which has been associated with increased chance of AMR and MDR acquisition. Our SDHSE MAGs demonstrated that MDR organisms (MDRO) are present in swine, and likely additional hosts contributing to global AMR. Altogether, our study provides comprehensive genetic support of MDR populations within the gut microbiome of swine.

## Introduction

Antimicrobial resistance (AMR) is a ubiquitous threat around the world, estimating to be the third cause of global human deaths^1^. Antimicrobial resistance was estimated to be associated with 4.95 million deaths globally in 2019^1^ and 2.8 million illnesses annually in the US alone^2^. AMR has also burdened medical systems and economies, and scientists see this as a sustained trend expecting $100-210 global losses due to AMR by 2050^3–5^. The burden and repercussions of AMR are a one world health concern as antibiotics are utilized for animals in addition to humans^6^. AMR describes bacteria containing genetic components which allow them to survive through antimicrobial treatment. Particular concern arises when bacteria exhibit resistance to multiple drugs^7,8^. These multidrug resistant (MDR) organisms (MDRO) can persist, at times, beyond all medicinally utilized antibiotics^7–9^. Moreover, multidrug resistant bacteria can spread through individuals, and between humans and animals, increasing the prevalence of AMR^10^. With the global burden of AMR, we need an enhanced understanding of AMR to combat infections caused by MDRO.

Animal agriculture has been identified as the largest antibiotic consumer^11^. Antibiotics have been used in agricultural animals, much like humans, to treat bacterial infections, but antibiotics are also for growth promotion in agriculture^12^. Swine production was estimated to be the current largest agricultural animal antibiotic-use sector in 2017^11,13^. Moreover, antimicrobial resistance rates are rising in the swine industry as the proportion of antibiotics, with resistance higher than 50%, increased in the swine industry from 0.13 to 0.34 from 2000 to 2018^14^. Global surveys^15–19^ and smaller-scale studies^20–26^ have in-large identified high consumption of tetracycline antibiotics in the past two decades in animal agriculture and swine. Tetracycline was estimated to account for 43% of antibiotic usage in agricultural animals from 2015 to 2017^19^. Unfortunately, tetracycline antibiotics are not exclusively utilized in animals. For example, chlortetracycline is used in both swine and humans^27,28^. With continued use of antimicrobial drugs, especially when utilizing the same treatments in humans and animals, and increasing resistance to antibiotics, AMR is a global concern to agriculture and humanity alike.

In monogastric animals, including pigs and humans, the gut microbiota has been identified as an AMR reservoir^29,30^. The gut microbiome has been recognized as a diverse environment in terms of antimicrobial resistant genes^31,32^. With oral antibiotic use, the gut has been demonstrated to increase in resistant bacteria^33^. Antibiotic treatments decrease the abundance of susceptible bacteria which allows resistant bacteria more resources, such as nutrients and space, to increase in abundance^34^. While the work in AMR is accumulating at a fast pace, we still have limited understanding on how genetic variations among the microbial populations contribute to the resistome.

Antibiotic use has been associated with bacteria containing increased genetic variation and so-called hypermutable bacteria^35,36^. Antibiotic usage selects for bacteria with genetic variation, or those with relatively high mutation rates termed hypermutable bacteria^35^. As bacteria develop variation, this leads to an increased chance of developing resistance^35,36^. Therefore with subsequent antimicrobial treatments, we are continually selecting for hypermutable populations harboring increased variation in turn having more opportunities for further AMR acquisition and MDR^35,36^. This can lead to MDR bacteria with high mutation rates to evade future antimicrobial treatments. However, studies related to the understanding of cumulative genetic variation across AMR genes in MDR bacteria in response to antibiotic supplementation (in vivo) among piglets are lacking. In studying microbial variation in these circumstances, we can further evaluate the risk of and potential treatments for MDR bacteria.

Clearly, we need a deeper understanding of antimicrobial resistance and MDR to enhance our approach to AMR. Here, we studied gut microbiota through two distinct antibiotic treatments (in-feed chlortetracycline [CTC] and in-feed tiamulin [TMU]) in addition to a non-antibiotic control (C). We utilized swine, with tetracycline and pleuromutilin class antibiotics, to provide an *in vivo* evaluation of a comparatively high and low utilized antibiotic classes^11,13^, in the global swine industry^15–26^. As mentioned previously, tetracycline antibiotics accounted for 43% of antibiotic usage in animal agriculture during 2015-2017 whereas pleuromutilin only accounted for 3%^19^. For our study, we performed metagenomic sequencing to obtain genes for functional analysis. Following our subsequent genome assembly and manual genome refining, we identified a subset of 11 metagenome-assembled genomes (MAGs) with high genetic variation prior to and throughout both antibiotic treatments and in control swine. We also confirmed consistent detection of the 11 MAGs and termed these MAGs: sustained detection and high sustained entropy (SDHSE) MAGs. Our SDHSE MAGs are of concern as they contained genetic variation and demonstrated MDR to both CTC and TMU. Moreover, we identified 22 distinct AMR genes in our SDHSE MAGs. Altogether, we provide evidence of MDR bacteria present in swine with concerningly high levels of genetic variation in 11 distinct microbial populations. Our research transcends global health with insights into antimicrobial resistance, and especially MDR, from a major contributor to global AMR.

## Materials and Methods

### Experimental design

The swine study was performed as previously described (Figure 1A-C)^37–39^. Swine (genetic line L337×1050, PIC, Hendersonville, TN) were housed in a commercial research nursery facility. Diets were fed with formulations as previously described^40^. All pigs were housed in one room with an enclosed, environmentally controlled, and mechanical ventilation system. Pens contained slatted floors with deep manure pits. Feed and water were provided *ad libitum* per pen with a six-hole stainless steel self-feeder (refilled via a robotic system) and pan waterer (Supplementary Table S1). This study utilized 648 pigs randomly distributed into 24 pens (27 pigs per pen), while working to minimize differences in average pen weight during distribution. Three treatments were administered, according to average pen weight, 7 days after weaning at 21 days of age, for a total of 14 days, each across 8 pens: control (no antibiotic; C), in-feed chlortetracycline (CTC; 22 mg/kg body weight; CTC-hydrochloride, Elanco Animal Health, Indianapolis, IN), and in-feed tiamulin (TMU; 5 mg/kg body weight; Denagard®, Elanco, Animal Health, Indianapolis, IN).

**Figure 1.**
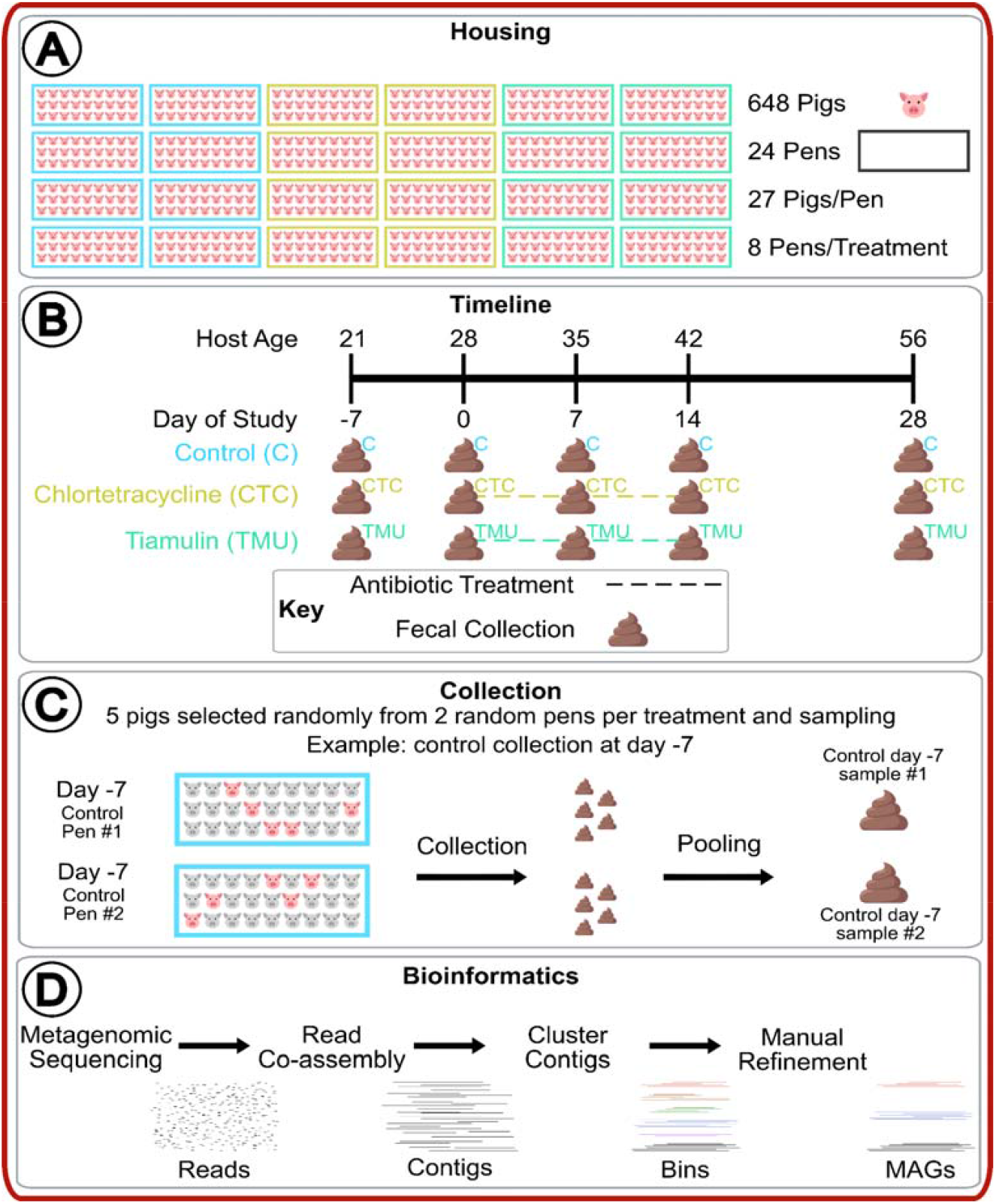
A) Pig and pen housing* allocation to treatments. B) Timeline of study. C) Fecal sample collection and pooling. D) Bioinformatics from sequencing reads to refined MAGs. *Image denotes pen treatments in same location for simplification. Note that pens were not all located in one region of room, instead pens were dispersed to control for adjoining pen interactions^37–39^.

Swine were managed according to protocol #4033 with Kansas State University Institutional Animal Care and Use Committee (IACUC). The authors also confirmed that all methods were performed in accordance with relevant guidelines and regulations^41^, and we affirmed that all methods were approved by Kansas State University.

### Sample Collection

For this study, we considered each pen as an experimental unit, and there were eight pens per antibiotic treatment (Figure 1A-C). Fecal collection occurred every seven days, starting on the day of introduction to the pens (Supplementary Table S1). Fecal samples were collected via gentle rectal massage from five randomly selected pigs per two random pens per treatment, and each fecal sample was stored in individual sterile plastic bags (Whirl-Pak® bags, Nasco, Ft. Atkinson, WI) and kept on ice during transportation. Processing occurred within 24 hours of collection, with intermittent storage at 4°C, at the Pre-harvest Food Safety laboratory, College of Veterinary Medicine, Kansas State University. Laboratory personnel were blinded to the treatments.

### DNA extraction

Fecal samples were stored at -80°C until DNA extraction. For each pen and time-point, the five fecal samples were pooled for DNA extraction (Figure 1A-C; n=30 samples [5 time-points*2 pens per treatment/time-point*3 treatments]). Total genomic DNA from fecal samples was extracted utilizing the DNeasy PowerSoil Pro Kits (QIAGEN Inc.; Valencia, CA), following the manufacturer protocols. We then quantified the extracted genomic DNA with a Nanodrop and Qubit™ (dsDNA BR Assay Kit [Thermo Fisher; Waltham, MA]) for DNA quality and concentration. Final storage of extracted DNA was at -80°C until library preparation and sequencing.

### Metagenomic sequencing and ‘omics workflow

Library preparation was performed on 30 samples with Nextera DNA Flex (Illumina, Inc.; San Diego, CA) (Figure 1B; Supplementary Table S1). A Tapestation 4200 (Agilent; Santa Clara, CA) was employed to visualize libraries followed by size-selected via a BluePippin (Sage Science; Beverly, MA). The final library pool of 30 samples was quantified with the Kapa Biosystems (Roche Sequencing; Pleasanton, CA) qPCR protocol, and sequenced on an Illumina NovaSeq S1 chip (Illumina, Inc.; San Diego, CA) with a 2 × 150 bp paired-end sequencing strategy.

We performed a bioinformatics workflow using anvi’o v.7.1 (https://anvio.org/install/; ‘anvi-run-workflow’ program)^42,43^. The workflow utilized Snakemake^44^ to perform multiple tasks: short-read quality filtering, assembly, gene calling, functional annotation, hidden Markov model search, metagenomic read-recruitment and binning^45^. Briefly, we processed sequencing reads using anvi’o’s ‘iu-filer-quality-minoche’ program removing low-quality reads following criteria described in Minoche *et al*.^46^. We termed the resulting quality-control reads “metagenome” per sample. We co-assembled quality-control short reads from metagenomes into longer contiguous sequences (contigs) according to no-treatment (prior to treatment/after) and treatment groups (C, CTC, TMU). We utilized MEGAHIT v1.2.9^42,47^ for co-assembly. The following anvi’o methods were then performed to further process contigs: (1) ‘anvi-gen-contigs-database’ to compute *k*-mer frequencies and identify open reading frames (ORFs) using Prodigal v2.6.3^42,48^; (2) ‘anvi-run-hmms’ to annotate bacterial and archaeal single-copy, core genes using HMMER v.3.2.1^42,49^; (3) ‘anvi-run-ncbi-cogs’ to annotate ORFs with NCBI’s Clusters of Orthologous Groups (COGs; https://www.ncbi.nlm.nih.gov/research/cog)^50^; and (4) ‘anvi-run-kegg-kofams’ to annotate ORFs from KOfam HMM databases of KEGG orthologs (https://www.genome.jp/kegg/)^51^.

We mapped all metagenomes’ short reads to contigs with Bowtie2 v2.3.5^52^. We converted mappings with samtools v1.9^42,53,54^ into BAM files. We profiled BAM mapping files (‘anvi-profile’)^42^ with a minimum length of 1,000 bp. We then combined profiles with ‘anvi-merge’ into a single anvi’o profile. Next, we used CONCOCT v1.1.0^55^ to group contigs into bins. We manually refined bins with ‘anvi-refine’ using bin tetranucleotide frequency and coverage across sample metagenomes^42,56,57^. After manual refining, we labeled bins that had ≥70% completion and <10% redundancy (both based on single-copy core gene annotation^58^) as metagenome-assembled genomes (MAGs). We analyzed MAG occurrences according to the “detection” metric. We determined single-nucleotide variants (SNVs) on all MAGs after read mapping with ‘anvi-gen-variability-profile’ and ‘--quince-mode’^42^. We used anvi’o’s DESMAN v2.1.1 to analyze SNVs to determine the number and distribution of subpopulations in the MAGs^59^. We accounted for non-specific mapping by removing any MAG subpopulations that made up less than 1% of the entire population and were explained by a singular MAG.

### Data analyses

We used RStudio v1.3.1093^60^ to visualize MAGs detection and entropy patterns in RStudio (https://www.rstudio.com/products/rstudio/) using: pheatmap (pretty heatmaps) v1.0.12^61^, ggplot2 v3.3.5 (https://ggplot2.tidyverse.org/)^62^, forcats v0.5.1 (https://forcats.tidyverse.org/)^63^, dplyr v1.0.8 (https://dplyr.tidyverse.org/)^64^, and ggpubr v0.4.0 (https://CRAN.R-project.org/package=ggpubr)^65^. We generated SNVs counts according to individual sample with anvi’o, anvi-summarize, and MAG entropies^66^ were generated with anvi’o’s anvi-gen-variability-profile^42,57^. Individual MAG entropy files and individual MAG statistical analysis files were combined respectively in RStudio with: tidyverse 1.3.1 (https://cran.r-project.org/web/packages/tidyverse/citation.html)^67^ and 1.4.0 (https://stringr.tidyverse.org/)^68^. We performed Welch two sample T-test^69^ statistical analysis on detection and entropy according to pre-treatment versus post-treatment and treatment groups. We used anvi’o COG annotations, as described above, for metabolic function analyses^42^. Our final figures were edited in Inkscape v1.2.1^70^.

### Data availability

We uploaded our metagenome raw sequencing data to the SRA under NCBI BioProject PRJNA899060. All other analyzed data, in the form of databases and fasta files, and bioinformatic scripts are accessible at figshare (https://doi.org/10.6084/m9.figshare.21548445.v1).

## Results and Discussion

Antimicrobial resistance (AMR), and especially multidrug resistance (MDR), are a global concern. Animal agriculture has been identified as the top consumer of antibiotics with the swine industry consuming the most of any agricultural sector^11,13^. In order to better understand AMR and MDR dynamics of the swine gut microbiome, we collected samples prior to, during and after antibiotic treatment. We utilized three distinct treatment groups: chlortetracycline, tiamulin, and non-antibiotic control. These antibiotics were utilized to allow analysis of distinctly utilized microbial classes across swine. Interestingly, we identified 11 distinct bacterial populations with similar detection levels pre- and post-treatment and between treatments. These bacteria harbored high genetic variation. The 11 microbial populations, assembled from our metagenomic data, were termed sustained detection and high sustained entropy (SDHSE) metagenome-assembled genomes (MAGs). Already exhibiting MDR, high variation in our resolved SDHSE MAGs could result in enhanced multidrug resistance. We further identified 22 unique AMR genes with varying detection in SDHSE MAGs. Altogether, we detailed AMR of swine microbiota with genetic support of existing MDR prior to antibiotic treatments and sustained variation throughout treatments. Our study advances AMR and MDR research by providing reflection on antibiotic and resistome association with animal agriculture, and potentially additional monogastric hosts.

### Resolved identify of gut metagenome-assembled genomes

We assembled and analyzed high resolution metagenome-assembled genomes (MAGs) to postulate functional distinctions between gut microbiota before and after antibiotic treatment. Each MAG represents a “microbial population.” We described a microbial population as an assemblage of coexisting microbial genomes in an environment that are similar enough to map metagenomic reads to the same reference genomes^71^. Metagenomic sequencing on an Illumina NovaSeq produced 772,688,506 paired-end reads from 30 fecal samples (Figure 1B; Supplementary Table S2). After quality filtering, 741,143,268 paired reads (96%) were utilized in contig co-assembly. We generated 330,769 contigs from assembly which described 1,018,536,193 nucleotides and 1,270,711 genes. We performed contig binning to create 369 bins, and after automatic and manual refinement we resolved 205 MAGs (Supplementary Table S3). To ensure high quality MAGs in our analysis, we performed downstream analysis with MAGs greater than 2M nucleotides (n=81), as these would more accurately represent bacterial genomes^72^. Of these 81 MAGs, each MAG, contained 360 ± 232 contigs and an N50 value of 18,345 ± 16,569 nucleotides. MAG GC contents ranged from 26% to 62%. Moreover, the average MAG size was 2,424,923 nucleotides. The MAGs were assigned to 6 bacterial phyla (*Actinobacteriota*, n=3; *Bacteroidota*, n=37; *Firmicutes*, n=38; *Planctomycetota*, n=1; *Proteobacteria*, n=1; and *Verrucomicrobiota*, n=1) with 96% of the MAGs resolved to 48 distinct genera.

We mapped each sample’s metagenomic reads (i.e. metagenome) to the 81 MAGs to determine detection throughout the study (Figure 2 and Supplementary Table S4). We confirmed detection of all 81 MAGs and determined general differential detection patterns according to detection clustering. The top branches broadly depict MAGs detected in the pre-treatment period. Comparatively, the middle clusters were sparsely detected. Finally, the bottom clusters were, in general, detected relatively high, compared to previous clusters, throughout the experiment regardless of pre- or post-treatment or treatment group. Altogether, our detection analysis suggested that association of microbial populations with swine hosts was far more complex than just what bacteria were affected by the use of antibiotics.

**Figure 2.**
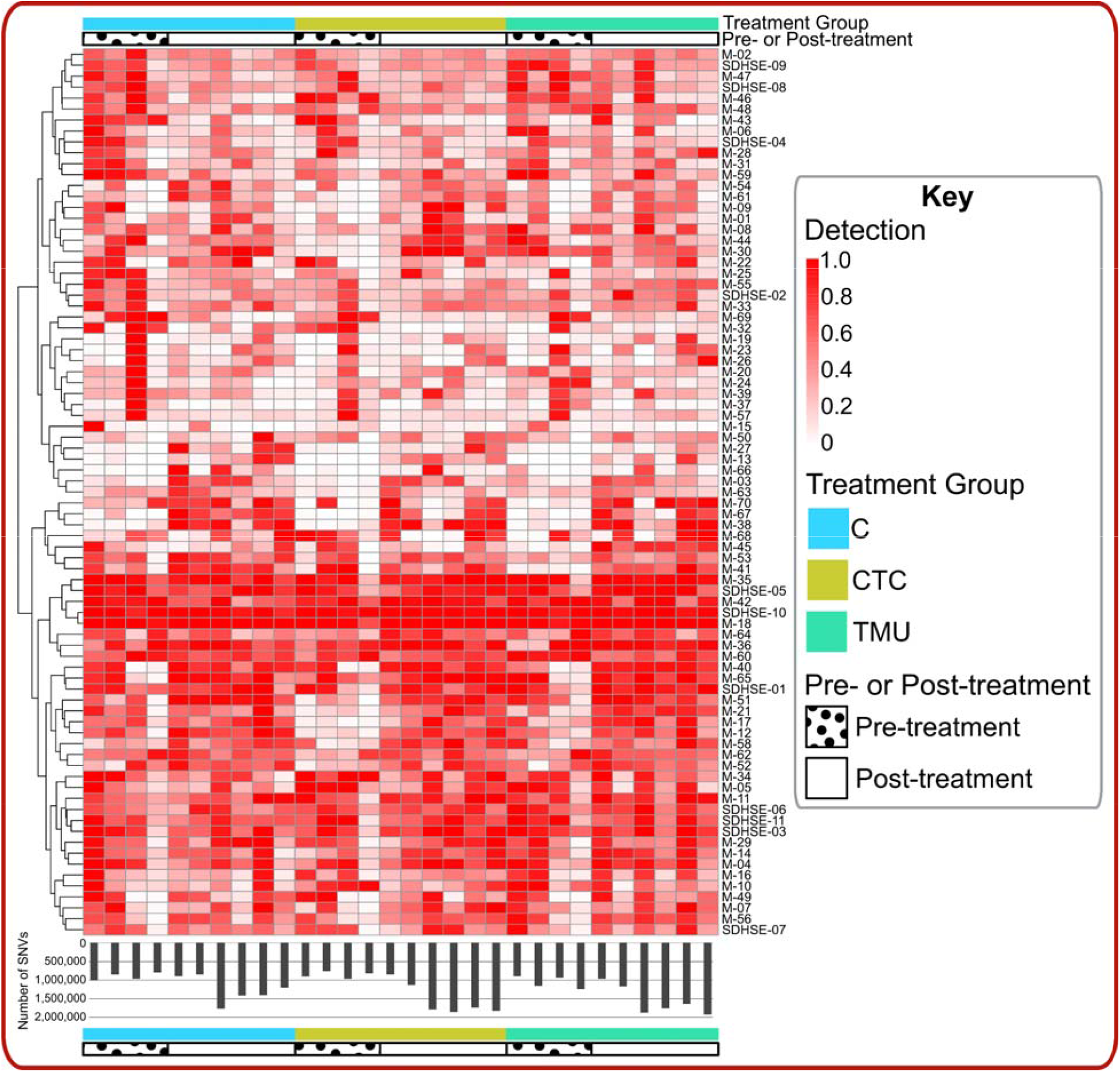
MAG detection heatmap (top) and single nucleotide variants (SNVs; bottom) according to treatment group and pre-/post-treatment (from left to right is earliest sampling [day -7] to last sampling [day 28] per treatment group) per sample.

Previous studies suggested environmental pressures, such as antibiotic administration, increased genetic variation in microorganisms^73,74^. The genetic variation in bacteria results from single nucleotide polymorphisms (SNPs), and could lead to generation of novel bacterial strains^73^. Studies further demonstrated that bacteria often used mutations as a mechanism for stress response, which is termed as stress-induced mutagenesis^75^. Since one of the mechanisms for the diversification and adaptation of the genomes operates at the single nucleotide level, we proceeded to resolve a more complete understanding of the environmental forces that affect adaptive strategies of our resolved MAGs to survive in the environment they resided in. Therefore, while our

MAGs were detected throughout the study, we were particularly interested in how MAG variants were changing according to treatment. Our bioinformatic analysis generated single nucleotide variants (SNVs) according to sample (Figure 2). We noticed relatively more SNVs associated with the post-treatment samples, which suggested that our resolved MAGs might respond to the antibiotic induced environmental pressure leading to the generation of new strains^73^. In light of this discovery, we proceeded to evaluate which MAGs were consistently high variation while maintaining detection even with different antibiotic treatments. These MAGs could potentially evade antimicrobial treatment with a multitude of variants, as demonstrated through sustained detection. Therefore, we next evaluated entropy throughout all 81 MAGs.

### MAGs harboring high genetic variation persisted through antimicrobial treatment

We performed single nucleotide variant (SNV) analysis to calculate entropy on our 81 MAGs to investigate genetic variation due antibiotic induced environmental pressure (Supplementary Table S5). Entropy describes nucleotide ratios for a given position, and entropy is measured from 0 (no variation; A=0, T=0, G=0, C=1) to 1 (complete variation; A=0.25, T=0.25, G=0.25, C=0.25)^76^. We performed statistical analysis to determine which MAGs held high sustained variation in the form of entropy and sustained detection (Supplementary Table S6). We discovered 31 MAGs with no statistical difference in entropy and detection (Supplementary Table S6). These MAGs represented microbial populations that were detected consistently regardless of antibiotic treatment. We further narrowed our selection to 11 MAGs with the relatively highest (33%) variation (Supplementary Table S6) because we were interested in MAGs harboring high variation, with potential multidrug evasion. Previous publications demonstrated the use of relative entropy analysis versus discrete entropy thresholds^77– 79^. These 11 MAGs were termed sustained detection and high sustained entropy (SDHSE) MAGs (Figure 3; Table 1). Of these SDHSE MAGs, 5 (45%) were assigned to the gram negative *Bacteroidota* (also known as *Bacteroidetes*)^80^ phylum, while 6 (55%) were annotated to gram positive *Firmicutes*. While members of both phyla have been associated with resistance to CTC and TMU, we identified only 2 (*Prevotella*^81–83^ and *Ruminococcus*^84–86^) of 9 genera associated with CTC resistance and 0 with TMU. Of our 11 SDHSE MAGs, 8 (73%) MAGs were annotated to 8 distinct species. Akin to the genus level, we provided novel associations of bacterial species, within the SDHSE MAG populations, exhibiting MDR. The finding indicated there are likely additional genera and species, with CTC and TMU resistance, then are currently known. Still, we wanted to investigate how the genetic variation of our SDHSE MAGs was related to AMR and MDR.

**Table 1.**
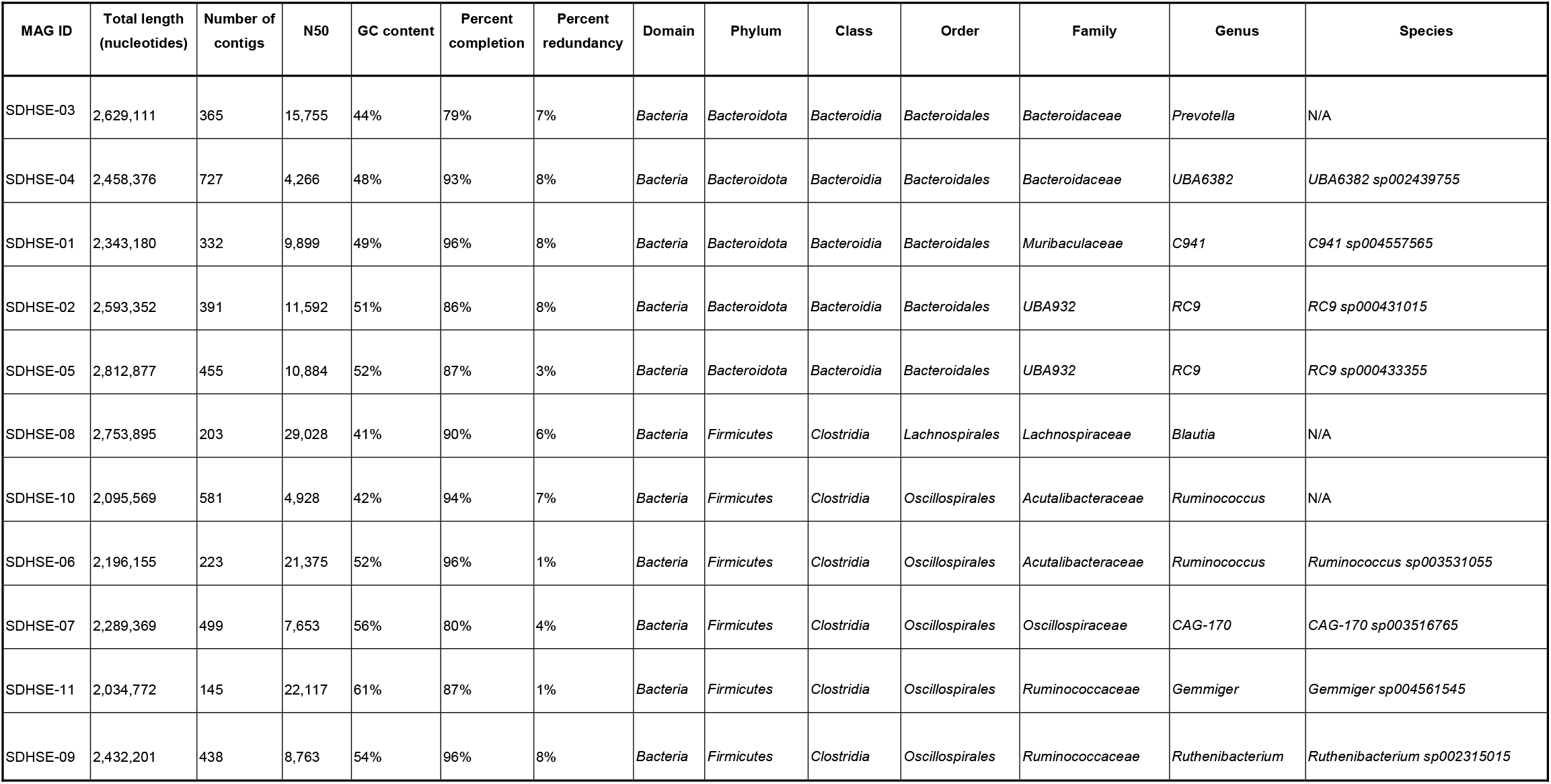
Taxonomic assignment and assembly statistics, of sustained detection and high sustained entropy (SDHSE) MAGs.

| MAG ID | Total length<br>(nucleotides) | Number of<br>contigs | N50 | GC content | Percent<br>completion | Percent<br>redundancy | Domain | Phylum | Class | Order | Family | Genus | Species |
| --- | --- | --- | --- | --- | --- | --- | --- | --- | --- | --- | --- | --- | --- |
| SDHSE-03 | 2,629,111 | 365 | 15,755 | 44% | 79% | 7% | Bacteria | Bacteroidota | Bacteroidia | Bacteroidales | Bacteroidaceae | Prevotella | N/A |
| SDHSE-04 | 2,458,376 | 727 | 4,266 | 48% | 93% | 8% | Bacteria | Bacteroidota | Bacteroidia | Bacteroidales | Bacteroidaceae | UBA6382 | UBA6382 sp002439755 |
| SDHSE-01 | 2,343,180 | 332 | 9,899 | 49% | 96% | 8% | Bacteria | Bacteroidota | Bacteroidia | Bacteroidales | Muribaculaceae | C941 | C941 sp004557565 |
| SDHSE-02 | 2,593,352 | 391 | 11,592 | 51% | 86% | 8% | Bacteria | Bacteroidota | Bacteroidia | Bacteroidales | UBA932 | RC9 | RC9 sp000431015 |
| SDHSE-05 | 2,812,877 | 455 | 10,884 | 52% | 87% | 3% | Bacteria | Bacteroidota | Bacteroidia | Bacteroidales | UBA932 | RC9 | RC9 sp000433355 |
| SDHSE-08 | 2,753,895 | 203 | 29,028 | 41% | 90% | 6% | Bacteria | Firmicutes | Clostridia | Lachnospirales | Lachnospiraceae | Blautia | N/A |
| SDHSE-10 | 2,095,569 | 581 | 4,928 | 42% | 94% | 7% | Bacteria | Firmicutes | Clostridia | Oscillospirales | Acutalibacteraceae | Ruminococcus | N/A |
| SDHSE-06 | 2,196,155 | 223 | 21,375 | 52% | 96% | 1% | Bacteria | Firmicutes | Clostridia | Oscillospirales | Acutalibacteraceae | Ruminococcus | Ruminococcus sp003531055 |
| SDHSE-07 | 2,289,369 | 499 | 7,653 | 56% | 80% | 4% | Bacteria | Firmicutes | Clostridia | Oscillospirales | Oscillospiraceae | CAG-170 | CAG-170 sp003516765 |
| SDHSE-11 | 2,034,772 | 145 | 22,117 | 61% | 87% | 1% | Bacteria | Firmicutes | Clostridia | Oscillospirales | Ruminococcaceae | Gemmiger | Gemmiger sp004561545 |
| SDHSE-09 | 2,432,201 | 438 | 8,763 | 54% | 96% | 8% | Bacteria | Firmicutes | Clostridia | Oscillospirales | Ruminococcaceae | Ruthenibacterium | Ruthenibacterium sp002315015 |

**Figure 3.**
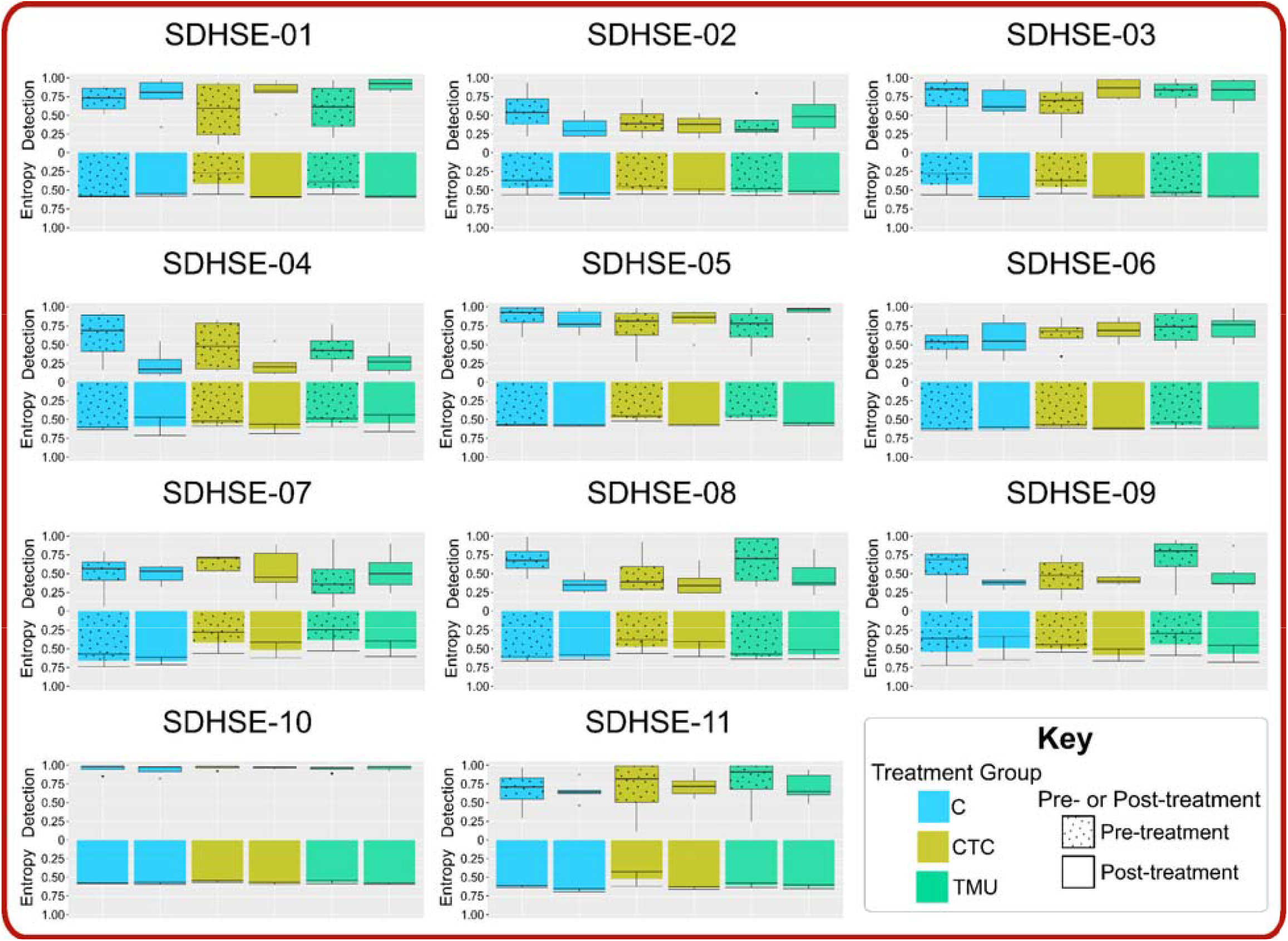
Detection boxplot and entropy bar graphs of our 11 sustained detection and high sustained entropy (SDHSE) MAGs.

Our SDHSE MAGs satisfied three important criteria - 1) consistent detection; 2) consistent high coverage of MAGs in the metagenomes; 3) consistent high variation of the MAGs in the metagenomes. Consistent detection demonstrated MDR, at least encompassing resistance to CTC and TMU, of the microbial populations. Consistent high coverage of the MAGs removed biases of identifying false variations among metagenomes due to coverage differences. Finally, previous publications have described how bacteria harboring variation are a concern for antimicrobial resistance^35,87,88^. When a microbial population contains a relatively high number of SNVs or contains a highly variable genetic background, the population contains genetic variation which may allow bacteria to persist even with antibiotic treatments. Here we demonstrated that our 11 SDHSE MAGs showed similar detection prior to and after distinct antibiotic treatments (CTC and TMU). The specific variants harbored in these MAGs are of particular interest to antimicrobial resistance (AMR) studies, thus, we surmised that the broad variation within these SDHSE MAGs likely contributed to the bacteria’s adaptive ability to survive antibiotic induced environments. Moreover, harboring continued high variation even after antibiotic treatment suggested many variants were able to persist during and after CTC and TMU treatment^35,87,88^. Previous studies highlight the role of antibiotic selection for populations with higher mutations, called hypermutable bacteria, which leads to high genetic variation in subsequent generations^35,36^. Our SDHSE MAGs were concerning as they contained high variation prior to antibiotic treatment and were able to remain present in the gut microbiome following antimicrobial treatment. Additional research is crucial to determine the presence of MDR organisms (MDRO) in additional hosts, regardless of previous antibiotic treatment. Our results suggested that there are likely numerous MDRO already present in hosts. Further antimicrobial treatments could continually be selecting for further MDR and hypermutable bacteria across all hosts, including across swine, monogastric and additional hosts. Hypermutable bacteria, including our SDHSE MAGs harboring numerous variants, are a concern to AMR with their MDR potential^35,36^.

We hypothesized that out 11 SDHSE MAGs likely contained AMR genes contributing to their continued detection. Therefore, we evaluated the MAGs for AMR genes within our functional potential annotations.

### Abundance of antimicrobial resistance (AMR) genes associated with sustained detection and high sustained entropy (SDHSE) MAGs

We hypothesized that genetic components associated with AMR supported the ability for SDHSE MAGs to prevail regardless of CTC and TMU use. We used COG annotations to investigate genetic functions for our 11 SDHSE MAGs, and we obtained a total of 21,025 COG annotations (average 1,911 per MAG). We observed numerous AMR genes within the high entropy contigs among the SDHSE MAGs (Supplementary Table S7). Within the COG annotations, we identified 19 unique gene annotations that coded for 18 distinct proteins or protein complexes related to AMR with an additional three genes (two complexes: YadH/YadG and RhaT) for drug efflux (Figure 4, Table 2, and Supplementary Table S8)^89–126^. We identified genes associated with six different drug efflux pump superfamilies (ATP-binding cassette [ABC], multidrug and toxic compound extrusion [MATE], drug/metabolite transporter [DMT], major facilitator [MFS], resistance-nodulation-division [RND], and small multidrug resistance [SMR]) alongside genes coding for: antimicrobial peptides (AMP), L-lactamases, L-lactamase regulators, and penicillin binding protein (PBP) relatives. Interestingly, of the 11 SDHSE MAGs, the gram negative MAGs (n=5) were, on average, annotated with 13 (57%) of the 22 genes, whereas the gram positive MAGs (n=6) were annotated on average with 12 (52%) genes. This agrees with previous literature indicating AMR is more often associated with gram negative bacteria relative to gram positive bacteria^127^. Still, both gram negative and gram positive bacteria cause significant illnesses and mortalities globally^127–129^. Given the risk MDR bacteria, including our SDHSE MAGs, pose to society, we further investigated individual resistance genes and proteins to build the knowledge surrounding AMR and MDR.

**Table 2.**
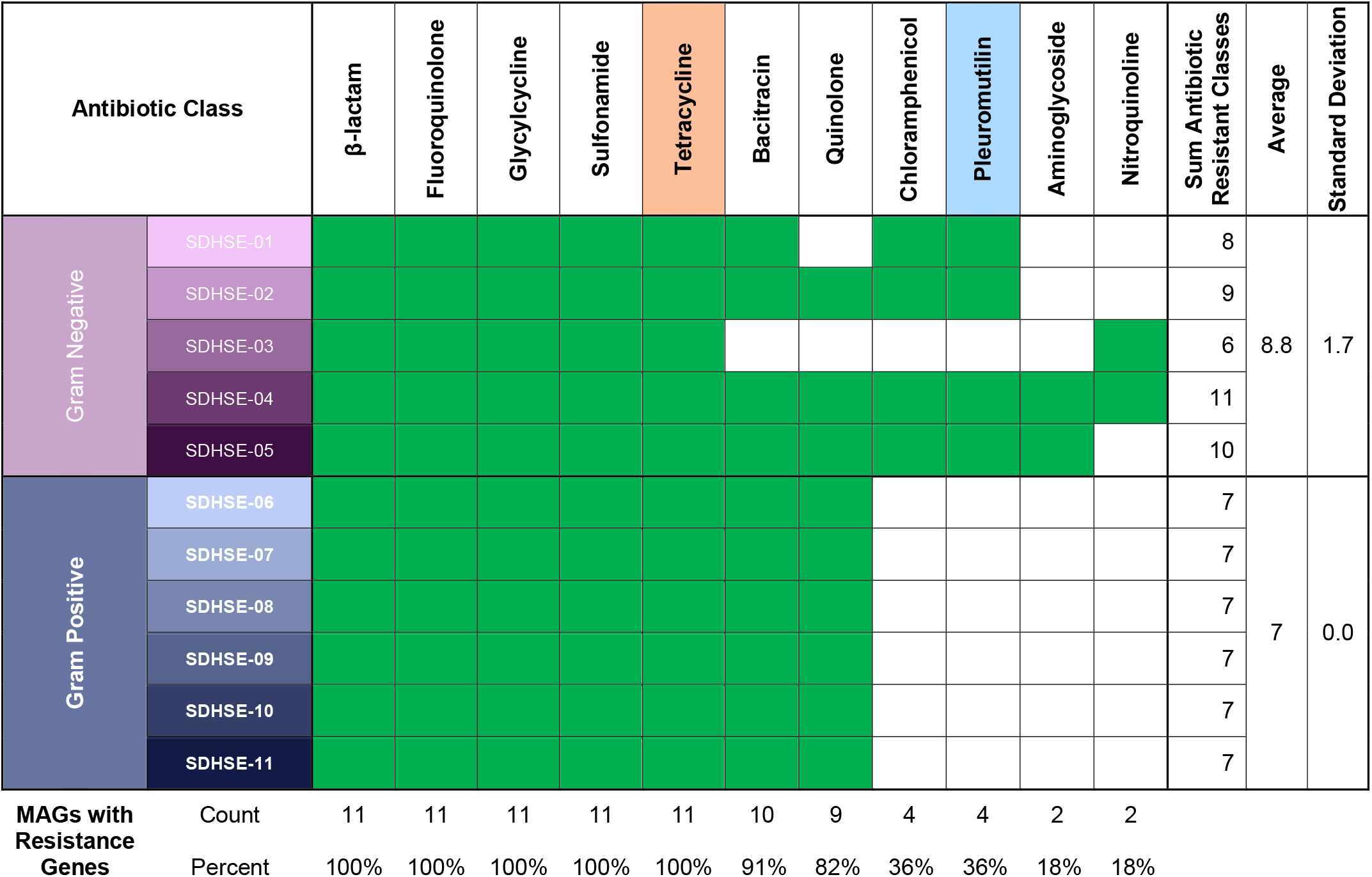
Antibiotic class resistance, based on previous publications and our annotated AMR and drug efflux genes, for our SDHSE MAGs^89–126^; green filled boxes indicate resistance associated with gene(s) whereas white demonstrates no AMR association.

**Figure 4.**
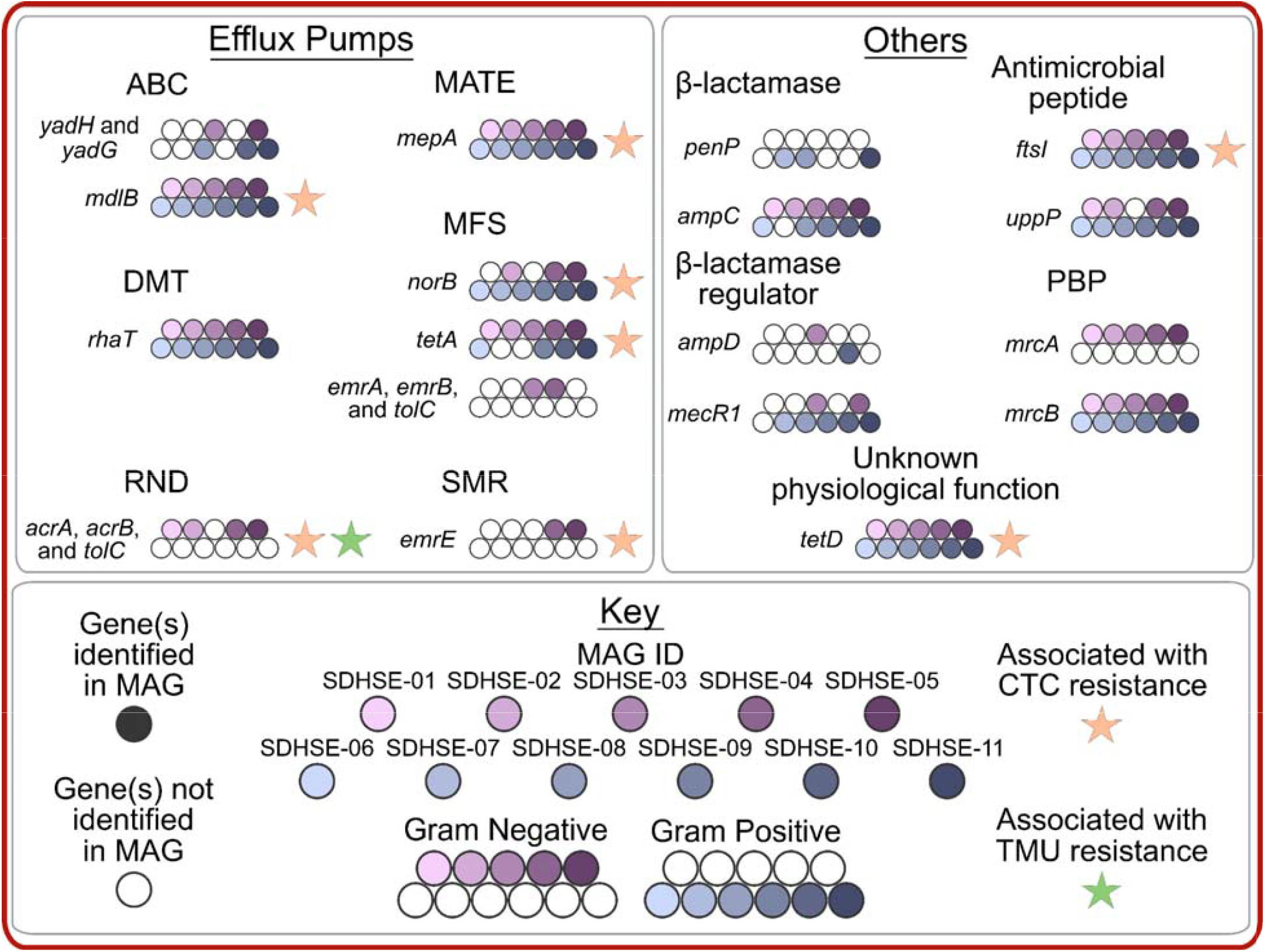
AMR genes detected in SDHSE MAGs annotated according to presence in our gram negative/positive MAGs and according to association with CTC (or tetracycline) or TMU (or pleuromutilin) previously^89–126^. Acronyms: ABC, ATP-binding cassette; MATE, multidrug and toxic compound extrusion; DMT, drug/metabolite transporter; MFS, major facilitator; RND, resistance-nodulation-division; SMR, small multidrug resistance; PBP, penicillin binding protein.

We noticed all SDHSE MAGs contained a variety of drug efflux pump and other (non-efflux pump) genes. Looking further into suspected resistance to antibiotics, based on AMR gene annotations, we discovered all SDHSE MAGs harbored AMR genes associated with 5 distinct antibiotic classes (Table 2)^89–115^. Tetracycline resistance, including resistance to CTC, is suspected across all SDHSE MAGs due to shared presence of *ftsI* and *mepA*, alongside *mdlB, norB, tetA, acrA*-*acrB*-*tolC, emrE*, and *tetD*^89–115^. The shared presence of multiple AMR genes could explain the consistent identification of these MAGs, regardless of antibiotic use. These bacteria could have repressed effects of the antibiotics as a result of these, and likely other, AMR genes. As expected, our gram negative MAGs contained, on average, a broader antibiotic class resistance (n=8.8) compared to gram positive MAGs (n=7.0)^127^. The physical membrane distinctions between gram positive and gram negative bacteria have resulted in greater antimicrobial resistance in gram negative bacteria^127^. Overall, we showed that all SDHSE MAGs demonstrated multidrug resistance potential which likely contributed to their continual presence even after antibiotic treatment.

We identified tripartite efflux pumps solely in gram negative MAGs. We identified the RND tripartite AcrAB-TolC complex genes in nearly of all our SDHSE gram negative genomes; only SDHSE-03 lacked AcrAB-TolC identification. RND pumps facilitate efflux across the outer membrane^124^. Gram positive bacteria lack this outer membrane which coincided with the absence of *acrA-acrB-tolC* annotation in our gram positive MAGs^124^. Moreover, since the majority of efflux pumps only facilitate efflux across the first membrane, RND pumps such as *acrA-acrB-tolC* have been described as broader and supports association of *acrA-acrB-tolC* with CTC and TMU resistance (Supplementary Table S8)^89,109,124^. A similar tripartite structure and efflux action have been described in the MFS efflux pump EmrAB-TolC, again in gram negative bacteria^124^. Although, EmrAB-TolC, to the best of our knowledge, has not been associated with CTC or TMU resistance. In our study, the identification of EmrAB-TolC in the CTC and TMU treatments suggested that EmrAB-TolC could have roles in CTC and TMU resistance. Further research is crucial to discover the range and action of tripartite efflux pumps in resistance and especially MDR.

We identified six of seven distinct efflux superfamilies, but we did not identify any proteobacterial antimicrobial compound efflux (PACE) annotations^130^. Although PACE has been demonstrated as prevalent in gram negative bacteria, previous literature has lacked identification in gram negative *Bacteroidota*, while PACE has been identified in gram positive *Firmicutes*^131^. PACE, the newest antibiotic class, was first described in 2015, with the second newest antibiotic class having been found in 2000^132^. Therefore, the breadth of knowledge surrounding PACE is growing and our MAGs could contain PACE efflux pump genes which have not been annotated within the COG database to date.

Beyond drug efflux pumps, we also identified genes coding for additional AMR proteins. As expected, we did not associate CTC or TMU resistance with L-lactamase or penicillin binding protein (PBP) related genes (Supplementary Table S8). L-lactamase inactivates L-lactam antibiotics, including penicillins, carbapenems, and cephalosporins^133^. We surmised that genes coding for these AMR proteins identified in our SDHSE MAGs have not previously been associated with CTC and TMU resistance due to their biochemical action, limiting their range of resistance^133^.

We did not identify any genes, outside of drug efflux genes, with suspected pleuromutilin (TMU) resistance. The only gene with suspected TMU resistance was the RND pump AcrAB-TolC^113^, which has also been associated with resistance of 5 other antibiotic classes. Only 36% of SDHSE MAGs contained genes with resistance to pleuromutilin antibiotics like TMU^109^. There are likely additional genes beyond the COG annotations we evaluated, and perhaps additional resistance which has yet to be discovered^134^. Additionally, we found that the majority of proteins (56%) associated with our genes had no previous support for resistance to CTC or TMU. Together, these results indicated a sizable knowledge gap in understanding the implications of strain level genetic variations among bacterial populations. Our SDHSE MAGs are likely harboring MDR to not only CTC and TMU, but other drugs, with genetic variations hindering targeted therapeutics^135^. While there has been a call for shifting our antibiotic usage to narrow or even strain-specific antibiotics to limit further AMR with application of broad antibiotics^136^, bacterial populations with high genetic variation could minimize the success of such therapies^35^. Clearly bacterial populations, such as SDHSE bacteria, with high genetic variation are concerning as they demonstrate increased AMR and threaten further AMR through targeted antimicrobials^35^. Future research needs to investigate similar SDHSE populations to determine their prevalence and risk they pose to global health.

## Conclusions

In order to investigate genetic variations pertaining MDR and AMR, we evaluated the gut microbiome for population dynamics before, during and after antimicrobial treatment. Our research is critical to understanding the implications of AMR on global health as we evaluated resistance in a sector dominating antibiotic use: swine production^11,13^. We demonstrated evidence of MDR bacterial populations present prior to antibiotic administration through 11 distinct bacterial populations we termed sustained detection and high sustained entropy (SDHSE) MAGs.

Within these MAGs, we indicated novel CTC and TMU resistance association with their taxonomic classifications at the genus and species levels. As work continues to discover gut-associated bacteria, we should evaluate their AMR characteristics to combat further resistance. Further highlighting the need for heightened AMR research, we found that approximately a third of our SDHSE MAGs contained annotated genes associated with TMU resistance. Although given the consistent identification of these MAGs during TMU treatment, there must be TMU resistance genes within the SDHSE genomes resulting in TMU resistance. Our SDHSE microbial populations harbored variation and AMR genes prior to antimicrobial treatment. We demonstrated that, although antimicrobial resistance is known to select for resistance^35,87,88^, resistant populations are currently present in the swine gut, indicating there are likely similar situations across additional hosts. While the number of antimicrobial resistance studies published has increased substantially since 2010^137^, the scientific community still has numerous topics to evaluate to better target AMR and MDR, all while under the pressure of rising antimicrobial resistance concerns^138^.

## Acknowledgements

Authors would like to thank the National Pork Board for funding part of this proposal (19-028). This work was supported in part by the USDA National Institute of Food and Agriculture, Hatch/Multistate Project 1014385. Our team is extremely grateful to the large number of individuals and organizations which assisted us in performing this research. We thank the University of Kansas Medical Center Genome Sequencing Facility for their expertise and assistance in sequencing including: Clark Bloomer, Dr. Veronica Cloud, Rosanne Skinner, and Yafen Niu. We greatly appreciate assistance from the following sources: Kansas State University Interdepartmental Genetics Program (fellowship for Brandi Feehan), Global Food Systems Seed Grant Program, Kansas Intellectual and Developmental Disabilities Research Center (NIH U54 HD 090216), the Molecular Regulation of Cell Development and Differentiation – COBRE (P30 GM122731-03) - the NIH S10 High-End Instrumentation Grant (NIH S10OD021743) and the Frontiers CTSA grant (UL1TR002366) at the University of Kansas Medical Center, Kansas City, KS 66160.

## Author Contributions

R. G. A, T. G. N, and M. D. T designed and executed the study at the commercial nursery facility. Sample collection was completed by R.G.A. R.G.A. performed DNA extraction while B.F and K.R. fulfilled Nanodrop and Qubit quality analysis. B.F. and Q.R., S.T.M.L. performed anvi’o bioinformatic analyses. B.F. and S.T.M.L. attributed biological relevance, wrote the manuscript, prepared figures, and supplementary files.

B.F. and S.T.M.L. performed major manuscript and figure refinement while remaining authors contributed to lighter refinement. All authors read, contributed to manuscript revision, and approved the submitted version.

## Supplementary Files

Supplementary Table S1. Demographics (diet, birth date, housing group, etc.) of swine hosts and dams, and sample metadata (swine age, host ID, stage and general health information, etc.).

Supplementary Table S2. Sequencing and assembly analysis including: metagenomic read counts initially obtained and assembly statistics according to co-assembly group.

Supplementary Table S3. Anvi’o results from initial bins and resulting MAGs, including taxonomic classification^42^.

Supplementary Table S4. Detection results from metagenome mapping to MAGs and SNV counts according to sample.

Supplementary Table S5. Entropy results from anvi’o.

Supplementary Table S6. Detection and entropy statistic results, and selection of SDHSE analysis.

Supplementary Table S7. COG annotations, including AMR annotations and AMR annotation summary, for SDHSE MAGs. Acronyms: ABC, ATP-binding cassette; MATE, multidrug and toxic compound extrusion; DMT, drug/metabolite transporter; MFS, major facilitator; RND, resistance-nodulation-division; SMR, small multidrug resistance; AMP, antimicrobial peptide; PBP, penicillin binding protein.

Supplementary Table S8. Antibiotic class resistance, based on previous publications, for our annotated AMR and drug efflux genes^89–126^; green filled boxes indicate resistance associated with gene(s) whereas white demonstrates no AMR association. Acronyms: ABC, ATP-binding cassette; MATE, multidrug and toxic compound extrusion; DMT, drug/metabolite transporter; MFS, major facilitator; RND, resistance-nodulation-division; SMR, small multidrug resistance; AMP, antimicrobial peptide; PBP, penicillin binding protein.

